# Roles for CEP170 in cilia function and dynein-2 assembly

**DOI:** 10.1101/2023.11.20.567836

**Authors:** Johannes F Weijman, Laura Vuolo, Caroline Shak, Anna Pugnetti, Aakash G Mukhopadhyay, Lorna R Hodgson, Kate J Heesom, Anthony J Roberts, David J Stephens

## Abstract

Primary cilia are essential eukaryotic organelles required for signalling and secretion. Dynein-2 is a microtubule-motor protein complex and is required for ciliogenesis via its role in facilitating retrograde intraflagellar transport from the cilia tip to the cell body. Dynein-2 must be assembled and loaded onto IFT-trains for entry into cilia for this process to occur but how dynein-2 is assembled at the base and how it is recycled back into a cilium remain poorly understood. Here, we identify Centrosomal Protein of 170 kDa (CEP170) as a dynein-2 interacting protein. We show that loss of CEP170 perturbs intraflagellar transport, Hedgehog signalling, and alters the stability of dynein-2 holoenzyme complex. Together, our data indicate a role for CEP170 in supporting cilia function and dynein-2 assembly.

**Summary:** Intraflagellar transport is required for the function of primary cilia. In this work, we show that Centrosomal Protein 170 (CEP170) interacts with the IFT motor dynein-2 and loss of CEP170 causes defects in dynein-2 assembly and cilia function.

## Introduction

Primary cilia (also called non-motile cilia) are hair-like organelles that protrude from nearly every vertebrate cell and are important for cell signalling. Mutations in cilia-related genes lead to a range of diseases, often developmental and skeletal, classified as ciliopathies (Huber and Cormier-Daire, 2012; Mill et al., 2023). Sonic hedgehog (Shh) signalling – one of the key pathways in vertebrate development – requires the primary cilium for proper signal transduction via Smoothened (Smo) (Corbit et al., 2005; Huangfu et al., 2003; Mill et al., 2023). Understanding how a cilium is built and regulated is critical in advancing our molecular understanding of ciliopathies.

At the core a cilium is the axoneme, a structure formed of doublet-microtubules with 9-fold radial symmetry (Hall and Hehnly, 2021). The cilium is separated from the cell body by the transition zone (TZ), a region of protein fibres that restrict protein entry into the cilium (Garcia-Gonzalo et al., 2011; Garcia-Gonzalo and Reiter, 2017). At the base of the primary cilium is the basal body originating from the mother centriole. The mother centriole, along with the daughter centriole and pericentriolar material, constitute the centrosome (Gonczy and Hatzopoulos, 2019; Tischer et al., 2021). Centrioles have a 9-triplet microtubule-based structure with 9-fold radial symmetry (Gonczy and Hatzopoulos, 2019; Tischer et al., 2021). The mother centriole is distinct from the daughter in that it has additional proteinaceous structures protruding radially from the distal end, called distal appendages (DAPs) and sub-distal appendages (sDAPs) (Hall and Hehnly, 2021). DAPs are required for cilia to form as, in non-dividing cells, ciliogenesis is initiated when pre-ciliary vesicles dock with DAPs, which eventually fuse with the plasma membrane forming the ciliary membrane (Ishikawa and Marshall, 2011; Tanos et al., 2013). DAPs form a barrier between the mother centriole and axoneme (Yang et al., 2018). There is conflicting evidence surrounding the role of sDAPs in ciliogenesis. Accumulated data support a model where sDAPs assemble a stabilized microtubule network that acts in ciliary vesicle docking ((Hehnly et al., 2012; Sorokin, 1962) reviewed in Hall and Hehnly (2021)).

The assembly and maintenance of a cilium requires intraflagellar transport (€FT) (Kozminski et al., 1993). IFT is an evolutionary conserved process by which material is transported in and out of the cilium, and is required to overcome the barrier of the TZ and concentrate cargoes within cilia (Lechtreck, 2015). IFT requires the assembly of polymeric trains composed of the protein complexes IFT-A and IFT-B, as well as the motor protein complexes, kinesin-2 and cytoplasmic dynein-2 (hereafter referred to as dynein-2) (Pigino, 2021; Webb et al., 2020). Anterograde trafficking is powered by kinesin-2, a heterotrimeric complex formed of the kinesin-family motor proteins, KIF3A and KIF3B, and kinesin-associated protein (KAP) (Webb et al., 2020). When the trains reach the tip of the cilium, the trains rearrange, and subsequent retrograde transport is driven by dynein-2. The correct assembly of IFT trains is vital to ensure proper ciliary composition and signalling. As well as receptors, signalling proteins, and cargo adaptors, kinesin-2 and dynein-2 motor proteins are themselves inactive cargoes during the directional transport step that they do not drive directly (Jordan et al., 2018; Toropova et al., 2017; Toropova et al., 2019).

Dynein-2 is a large (>2 Mda) multisubunit complex responsible for minus-end directed IFT. In humans, dynein-2 is composed of dynein-2 heavy chain (DYNC2H1), one copy each of the intermediate chains, WDR60 and WDR34, the light intermediate chain LIC3 (DYNC2L1), and four light chains, Roadblock-1 and -2 (DYBLRB1 and DYNLRB2), LC8-1 and LC8-2 (DYNLL1 and DYNLL2), TCTEX-1 (DYNLT1), TCTEX-3 (DYNLT3), all of which are also found in dynein-1, and TCTEX1D2 (DYNLT2B) (Asante et al., 2014; Toropova et al., 2019; Vuolo et al., 2020). Mutations in all dynein-2 subunits have been shown to cause ciliopathies, often leading to defects in bone development such as Jeune asphyxiating thoracic dystrophy (JATD) (also known as short-rib thoracic dysplasia (SRTD)) (Huber and Cormier-Daire, 2012; Mill et al., 2023).

Previous work has defined interactions of WDR60 and WDR34 with other dynein-2 components and IFT proteins (Hiyamizu et al., 2023b; Shak et al., 2023; Vuolo et al., 2018). In this study, we extend that work to define the sDAP protein CEP170 as a dynein-2 interactor. We show that dynein-2 interacts with CEP170 and that cells lacking CEP170 inefficiently assemble the dynein-2 holoenzyme as well as having minor ciliary defects. Our data suggests a role for CEP170 in supporting cilia homeostasis as well as dynein-2 assembly.

## Results

### CEP170, but not CEP170B, interacts with dynein-2

Previously, we have used proteomic data generated using tagged WDR60 and WDR34 (Hiyamizu et al., 2023b; Shak et al., 2023; Vuolo et al., 2018) to identify dynein-2 interacting proteins. The most consistent hit in these data sets, CEP170, was identified in 12 new and previously published tandem mass tag (TMT) datasets, 11 of which having a Log_2_ abundance ratio greater than 1 (Fig. 1A, Table S1). By comparison, the related protein, CEP170B was only found in 5 of those datasets, and only in 2 with a Log_2_ abundance ratio of greater than 1 (Fig. 1A, Table S1). We also previously identified CEP170 binding in non-TMT proteomic methods (Table S1) (Hiyamizu et al., 2023b). We validated the binding of CEP170 to WDR34 by coimmunoprecipitation (Fig. 1B). CEP170B has 33.4% identity with CEP170 and similar centrosomal localisation to CEP170 (Fig. S1). However, we could not detect CEP170B in co-immunoprecipitation assays with HA-WDR34 (Fig. 1C) suggesting that the interaction is specific to CEP170. Interestingly, immunoprecipitation with HA-WDR34 in the background of *WDR60* KO cells (Vuolo et al., 2018) led to the isolation of more CEP170 (Fig. 1B). The fact that we reliably identify CEP170 in experiments with both WDR34 and WDR60 in these experiments strongly suggests an interaction with the dynein-2 holocomplex.

**Figure 1:**
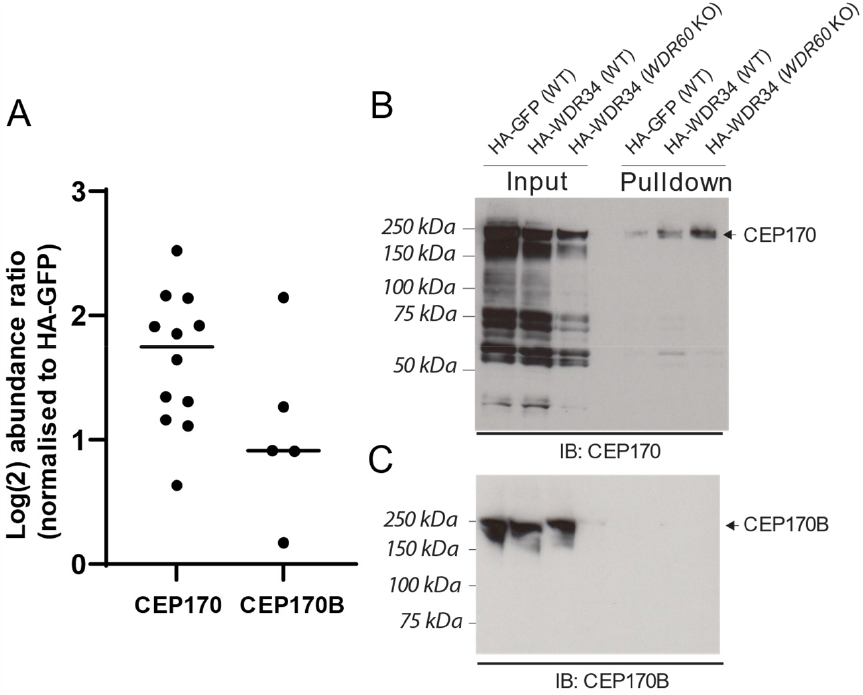
Dynein-2 interacts with CEP170 but not CEP170B. (A) Previous data sets from either GFP-WDR34, HA-WDR34 or HA-WDR60 interaction tandem mass tag (TMT) proteomics have reliably identified CEP170 being pulled-down with a Log_2_ abundance ratio above 1. Data is presented as Log_2_ abundance ratio, normalized to HA-GFP expression. A breakdown is shown in Table S1. Lines represent mean. Co-immunoprecipitation of CEP170 (B) but not CEP170B (C), in WT RPE1 cells expressing HA-WDR34. When performed in *WDR60* KO cells more CEP170 is pulled down.

CEP170 is a centrosomal protein, known to locate to sDAPs on mature centrioles (Guarguaglini et al., 2005). We therefore sought to investigate whether CEP170 may have a role in cilia function.

### CEP170 KO cells can still form cilia

To further study the role of CEP170 in ciliogenesis and cilia function, we generated *CEP170* KO RPE1 and mouse IMCD3 cells (Fig. S2) and performed serum-starvation induced ciliation assays (Fig. 2). *CEP170* KO RPE1 cells were able to extend cilia at the same proportion as WT RPE1 cells (Fig. 2A and B). In one of the RPE1 *CEP170* KO clones (23G7) there was a modest, but consistent, increase in cilia length (Fig. 2C). In IMCD3 cells, *CEP170* KO cells were still able to ciliate (Fig. 2D) and the cilia were of comparable lengths to WT cells (Fig. 2E). However, we did observe that *CEP170* KO IMCD3 cells ciliated at a reduced proportion compared to WT IMCD3 cells (Fig. 2F).

**Figure 2:**
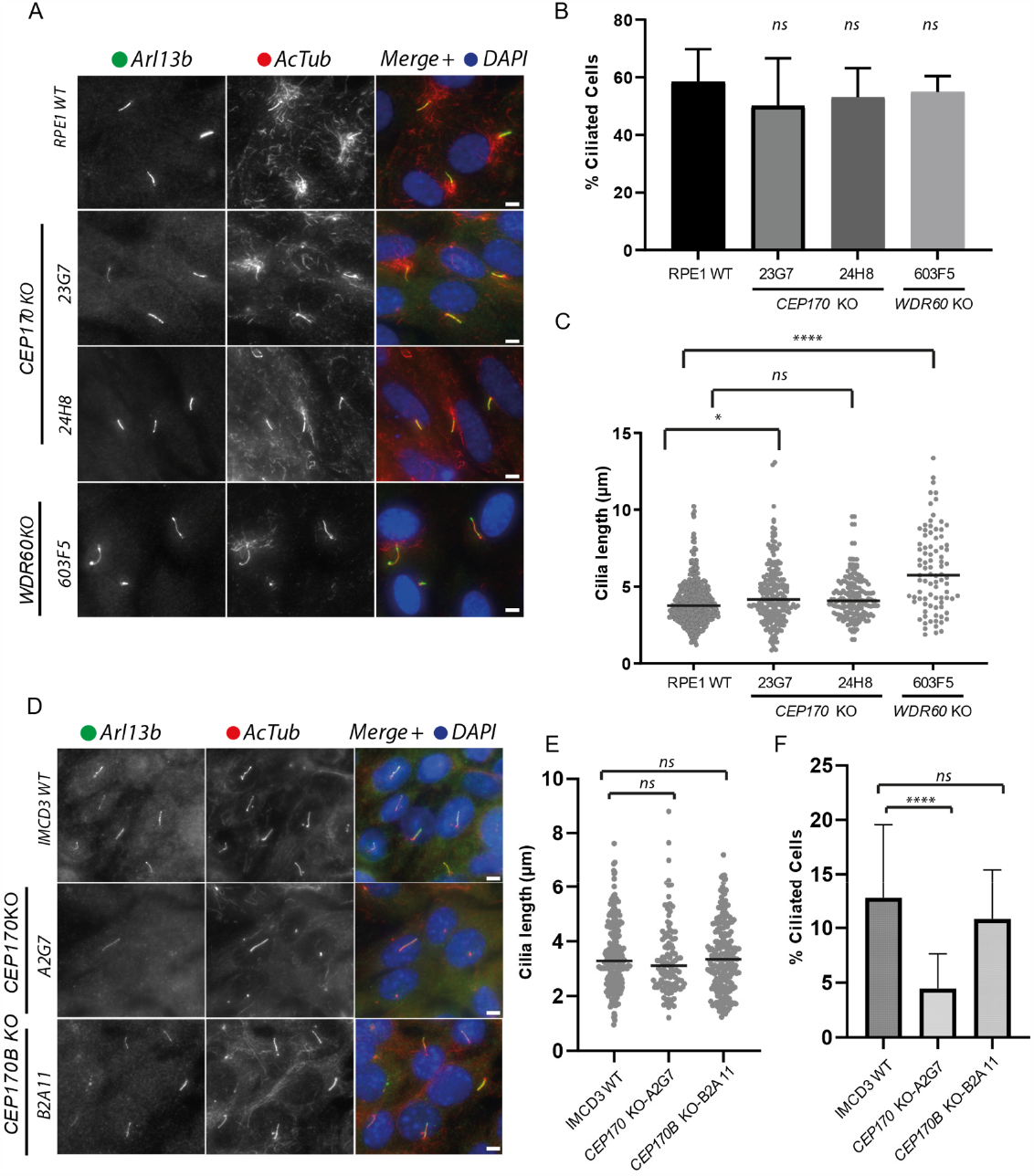
A role of CEP170 in ciliogenesis in RPE1 and IMCD3 cells. (A) Cilia labelled to detect Arl13b (green) and acetylated tubulin (AcTub, red) in RPE1 WT, *CEP170* KO (clones 23G7 and 24H8) and *WDR60* KO (clone 603F5) cell lines. (B) Cilium length in *CEP170* KO (23G7 and 24H8) and *WDR60* KO (603F5) compared with WT cells (n = 3; 367 WT, 212 *CEP170* KO (23G7), 174 *CEP170* KO (24H8), 87 *WDR60* KO (603F5) cells quantified). Plotted is mean with SD. (C) Percentage of ciliated cells (n = 3; 636 WT, 429 *CEP170* KO (23G7), 139 *CEP170* KO (24H8), 159 *WDR60* KO (603F5) cells quantified). Line represents median. One-way ANOVA followed by Kruskal-Wallis test, * p=0.022, **** p < 0.0001, ns – not-significant. (D) Cilia were stained with the markers Arl13b (green) and acetylated tubulin (AcTub, red) in IMCD3, *CEP170* KO and *CEP170B* KO cell lines. (E) Cilium length in *CEP170* KO (clone A2G7) and *CEP170B* KO (clone B2A11) compared with WT cells (n = 4; 213 WT, 112 *CEP170* KO (A2G7), 218 *CEP170B* KO (B2A11) cilia quantified). Line represents median. (F) Percentage of ciliated cells (n = 4; 1607 WT, 2661 *CEP170* KO (A2G7), 2208 *CEP170B* KO (B2A11) cells quantified). Plotted is mean with SD. One-way ANOVA followed by Kruskal-Wallis test, **** – p < 0.0001, ns – notsignificant. Scale bars = 5 μm.

CEP170 is recruited to sDAPs by ninein (Graser et al., 2007). In ciliated cells, ninein was still recruited to the ciliary basal body (Fig. S3A), indicating that our *CEP170* KO and *WDR60* KO cells still have sDAPs. We were able to further confirm this by electron microscopy (EM) on non-ciliated cells (Fig. S3B).

The ciliary TZ zone is an important barrier that separates the axoneme of the cilium to the rest of the cytoplasm, and disruption to the TZ is associated with numerous ciliopathies (Garcia-Gonzalo et al., 2011; Garcia-Gonzalo and Reiter, 2017). In *CEP170* KO cells the TZ proteins RPGRIP1L and TMEM67 were both localized correctly indicating that the TZ is not grossly impacted by loss of CEP170 (Fig. S4).

Previous work has suggested a role for CEP170 in cilia disassembly (Lamla, 2009). We tested this by readdition of FBS to serum starved cells. *CEP170* KO cells remained ciliated for a much longer period than WT cells or cells lacking WDR60 (Fig. S5A and B). Live cell imaging showed that once detected, cilia excision proceeds similarly in both WT and *CEP170* KO cells i.e. with considerable variability in timing but visually the same with respect to GFP-Arl13B (Fig. S5C).

### IFT88 accumulates at the ciliary tip in CEP170 KO cells

We next sought to address whether loss of CEP170 had any effect on IFT. IFT88 is a component of IFT-B and accumulations in ciliary tips can be used to infer defects in IFT (Hou and Witman, 2015). Serum-starved cells were fixed and labelled to detect IFT88. In *CEP170* KO RPE1 cells, we saw an increase in the proportion of cells that had IFT88 accumulations at the ciliary tip, similar to what we see in *WDR60* KO cells (Fig. 3A and B). In IMCD3 cells, *CEP170* KO, but not *CEP170B* KO, caused an increase in the relative ciliary tip fraction of IFT88 whereas there was no difference in the relative fraction at the base (Fig. 3C-E). These data indicate that loss of CEP170 could lead to mild defects in IFT leading to an accumulation of IFT88 at the ciliary tip.

**Figure 3:**
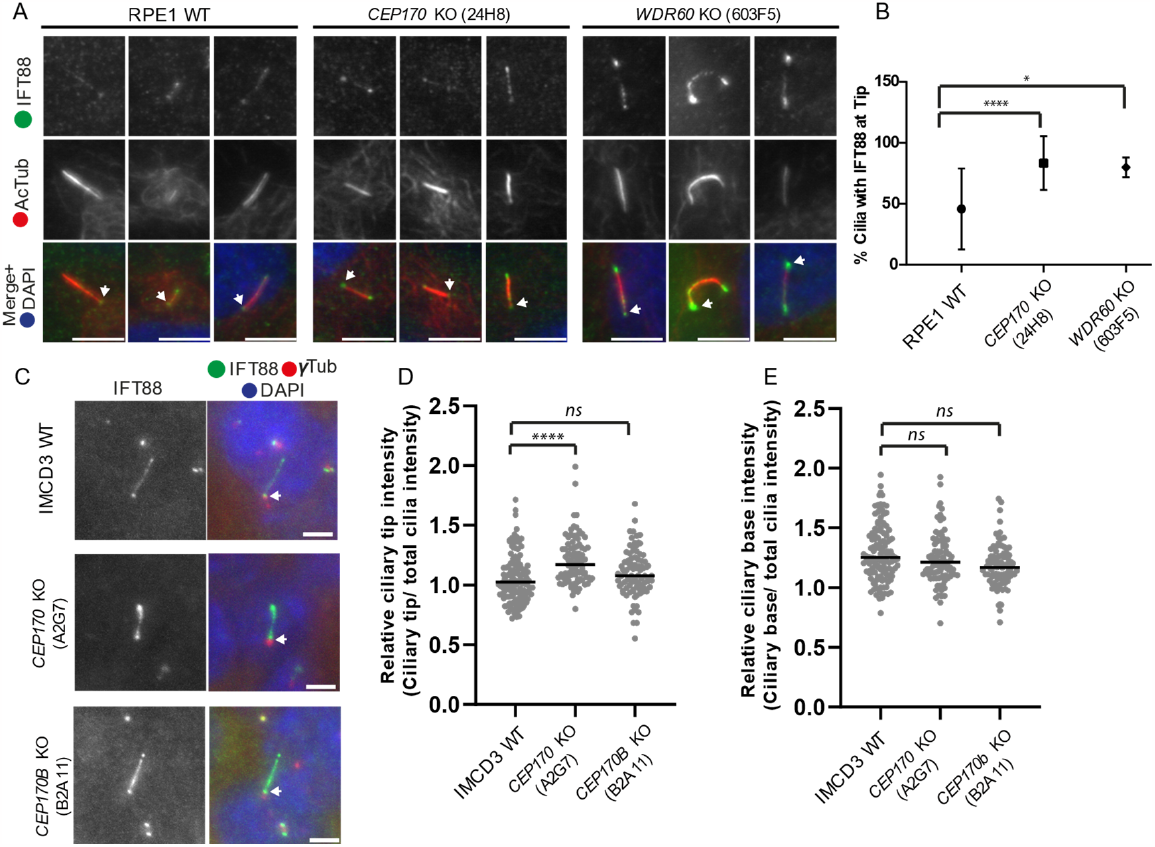
Localization of IFT88 in CEP170 KO cilia. (A) Localization of IFT88 (green) in cilia (acetylated tubulin, AcTub, red) in RPE1 WT, *CEP170* KO (clone 24H8) and *WDR60* KO (clone 603F5) cell lines. Scale bars = 5 μm. (B) Percentage of cilia showing IFT88 staining at cilia tips (n = 3; 167 WT, 137 *CEP170* KO (24H8), 38 *WDR60* KO (clone 603F5) cells quantified, one-way ANOVA followed by Kruskal-Wallis test, * – p=0.0191, **** – p < 0.0001). Plotted is mean with SD. (C) Localization of IFT88 (green) in cilia in IMCD3 WT, *CEP170* KO (clone A2G7) and *CEP170B* KO (clone B2A11) cell lines. Scale bars = 5 μm. The basal body is marked with gamma tubulin (γTub, red). (D and E) ImageJ plot profile tool was used to quantify the IFT88 intensity at the tip (D) and at the base (E). n=3; cells quantified: 213 IMCD3, 112 *CEP170* KO (A2G7), 218 *CEP170B* KO (B2A11). One-way ANOVA followed by Kruskal-Wallis test, ns – not significant, **** – p < 0.0001. Bars represent means. Arrows indicate basal body.

### CEP170 KO cells do not display major defects in IFT dynamics

To further examine the IFT88-tip accumulations, we generated a *CEP170* KO (Fig. S4) in IMCD3-*FlpIn*-*IFT88-NeonGreen x3* (NG3) cells (Liew et al., 2014; Mukhopadhyay et al., 2010; Ye et al., 2018) to allow us to follow IFT in real-time using total internal reflection fluorescence (TIRF) microscopy. Live imaging (Movies 1-3) and kymograph analysis (Fig. 4A and B) was used to measure anterograde and retrograde velocities and events (Fig. 4C-F). We observed a slight increase in anterograde velocity for one of *CEP170* KO clone (Fig. 4D) but there was no change in retrograde velocity (Fig. 4C) nor the number of events for transport in either direction (Fig. 4E and F).

**Figure 4:**
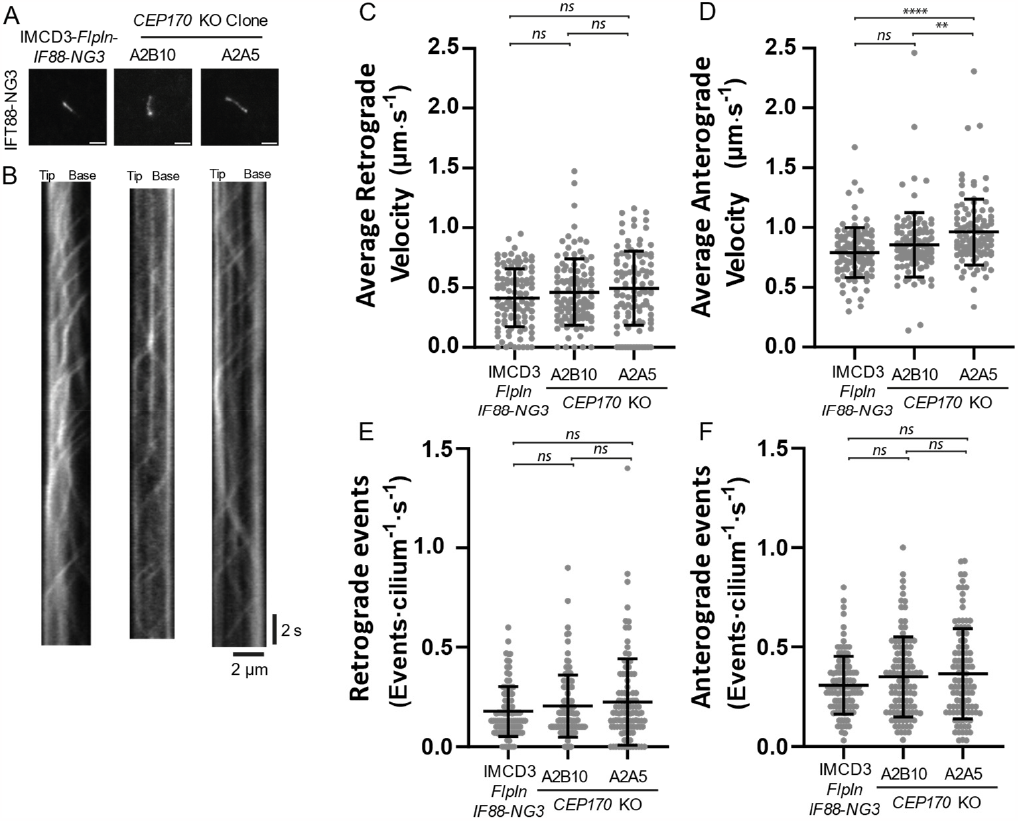
Live TIRF imaging of IFT88-NG in CEP170 KO cells. (A) IMCD3-*FlpIn*-*IFT88-NG3* and *CEP170* KO (clones A2B10 and A2B5) cell lines were serum starved and cilia were imaged live by TIRF microscopy (example movies are shown in Movies 1-3). Scale bars = 2 μm. (B) From individual cilia, kymographs were generated and used to measure retrograde (C) and anterograde (D) velocities and calculate retrograde (E) and anterograde (F) events. N=3, cells analysed: 105 WT, 107 *CEP170* KO (clone A2B10), 104 (clone A2A5). Bars represent mean ± standard deviation. One-way ANOVA followed by Kruskal-Wallis test, ns – not significant, ** – p=0.0019, **** – p < 0.0001.

### Heightened Smo response in CEP170 KO cilia

Sonic hedgehog (Shh) signalling requires functional cilia (Breslow et al., 2018). In basal conditions, the membrane receptor Smoothened (Smo), part of the Shh pathway, localizes to cilia but is rapidly exported by a process involving retrograde IFT. When Shh signalling is stimulated, by Shh or an agonist such as Smoothened agonist (SAG), Smo instead accumulates within the cilium (Corbit et al., 2005; Hamada et al., 2018; Vuolo et al., 2018). In absence of SAG, Smo is largely absent from cilia in WT and *CEP170* KO cells (Fig. 5A and C). In the presence of SAG, we found that a greater proportion of cilia in *CEP170* KO cells accumulated Smo than in WT cells (Fig. 5B and C).

**Figure 5:**
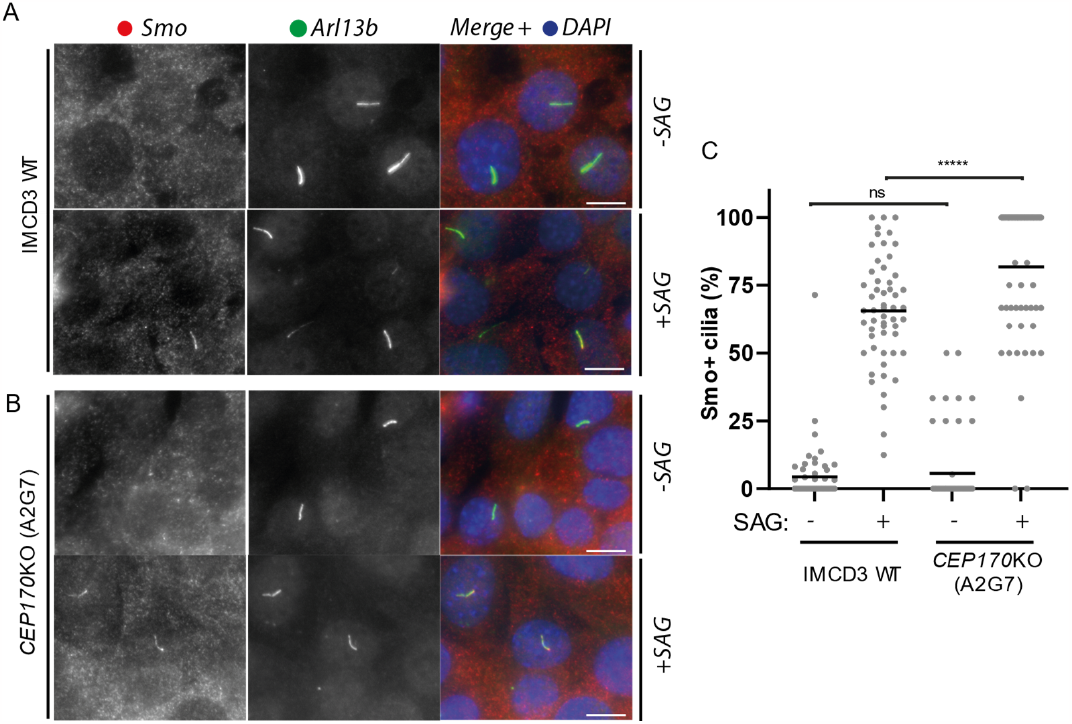
SMO response in *CEP170* KO Cells. (A and B) IMCD3 WT (A) or *CEP170* KO (clone A2G7) (B) cells were serum starved in the absence or presence of SAG. Cells were fixed and stained for Smo (red) and the cilia marker Arl13b (green). Scale bars = 10 μm. (C) Quantification of percentage Smo positive cilia of indicated IMCD3 cell lines in presence or absence of SAG. N=3, cells counted: IMCD3 WT 408 -SAG, 513 +SAG; *CEP170* KO (A2G7) 123 -SAG, 142 +SAG. Mann-Whitney test was Used. ns – not significant, **** – p<0.0001. Bars represent means.

### Dynein-2 holocomplex assembly is disrupted in CEP170 KO cells

Mutations in dynein-2 have been linked to Shh dysregulation (May et al., 2005). Therefore, we sought to examine the localization of the dynein-2 heavy chain (DHC2) directly. Notably, immunofluorescence showed a decrease in DHC2 localization to cilia in *CEP170* KO cells compared to WT (Fig. 6A-C). We also noted a decrease in DHC2 labelling at the base of the cilium (Fig. 6D-E). No differences in expression of DHC2 or other dynein-2 subunits were observed by immunoblot (Fig. 6F), suggesting that the reduction in ciliary and basal body DHC2 does not relate to changes in dynein-2 overall expression. As we have done previously (Vuolo et al., 2018), we used proteomic profiling to allow us to look at dynein-2 assembly (Fig. 7). In WT RPE1 cells, using HA-tagged WDR60, we could detect DHC2, the intermediate chain WDR34, dynein-2 light chains and individual components of IFT-B as well as the IFT-A component, IFT140 (Fig 7A). This latest dataset is consistent with our earlier findings that CEP170, but not CEP170B, is reliably co-immunoprecipitated with HA-WDR60 (Fig 7A and Table S1). We repeated the analysis in *CEP170* KO cells (Fig. 7B). We could detect DHC2, WDR34 and light chains as well as the chaperone NUDCD1 (known to be important in WD-repeat protein folding (Asante et al., 2014; Taipale et al., 2014; Vuolo et al., 2018)p. Other than the light chain proteins, DYNLT1 (TCTEX1) and DYNLT3 (TCTEX3) (which are also known subunits of dynein-1 (Vuolo et al., 2018; Vuolo et al., 2020)), all identified WDR60 binding partners had a Log_2_ abundance ratio below 0. These data strongly suggest a role for CEP170 in the assembly or stabilization of the dynein-2 holocomplex.

**Figure 6:**
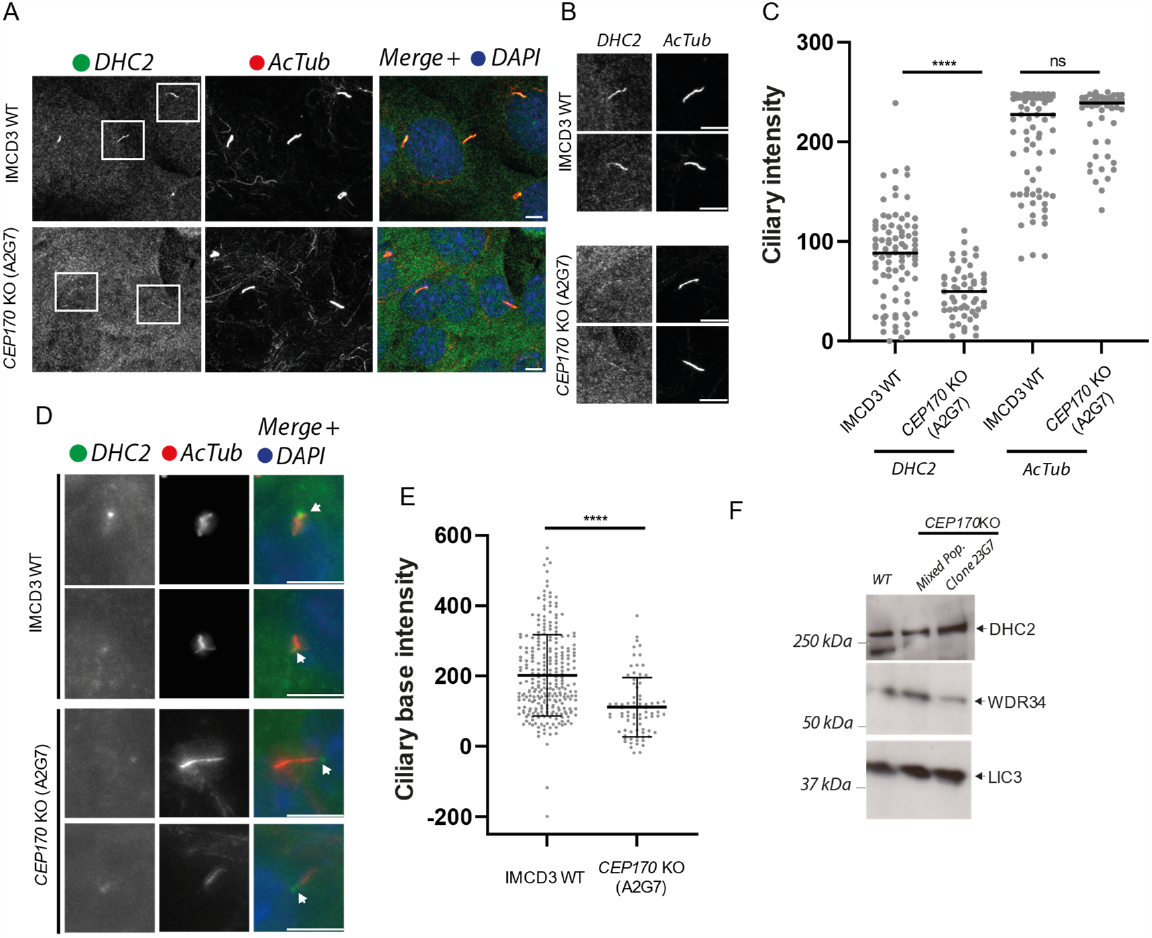
Dynein-2 heavy chain cilia localization in CEP170 KO cells. (A) IMCD3 WT and *CEP170* KO (clone A2G7) cells were serum starved, fixed, and labelled for DHC2 (green) acetylated tubulin (AcTub, red) (B) zoom of boxed cilia from (A). Scale bars = 5 μm. (C) Intensity of the full length of the cilia was quantified. As a control, AcTub intensity was also quantified and intensities were similar in WT and KO cells (n = 3; 85 IMCD3 WT, 55 *CEP170* KO (A2G7) cells quantified. Mann-Whitney test was used, **** p <0.0001. (D) Serum starved cells were fixed and immunolabelled to detect DHC2 (green) and AcTub (red). Arrows indicate the basal body. Scale bars = 5 μm. (E) Basal body DHC2 staining intensity (background normalized) comparing WT and *CEP170* KO (A2G7) IMCD3 cells. N = 3; 269 IMCD3 WT, 80 *CEP170* KO (A2G7) cells quantified). Mann-Whitney test was used, **** – p <0.0001. (F) Immunoblot of lysates from either WT or *CEP170* KO (mixed population (pop) or clone 23G7) cells showing indicated dynein-2 subunits are still expressed.

**Figure 7:**
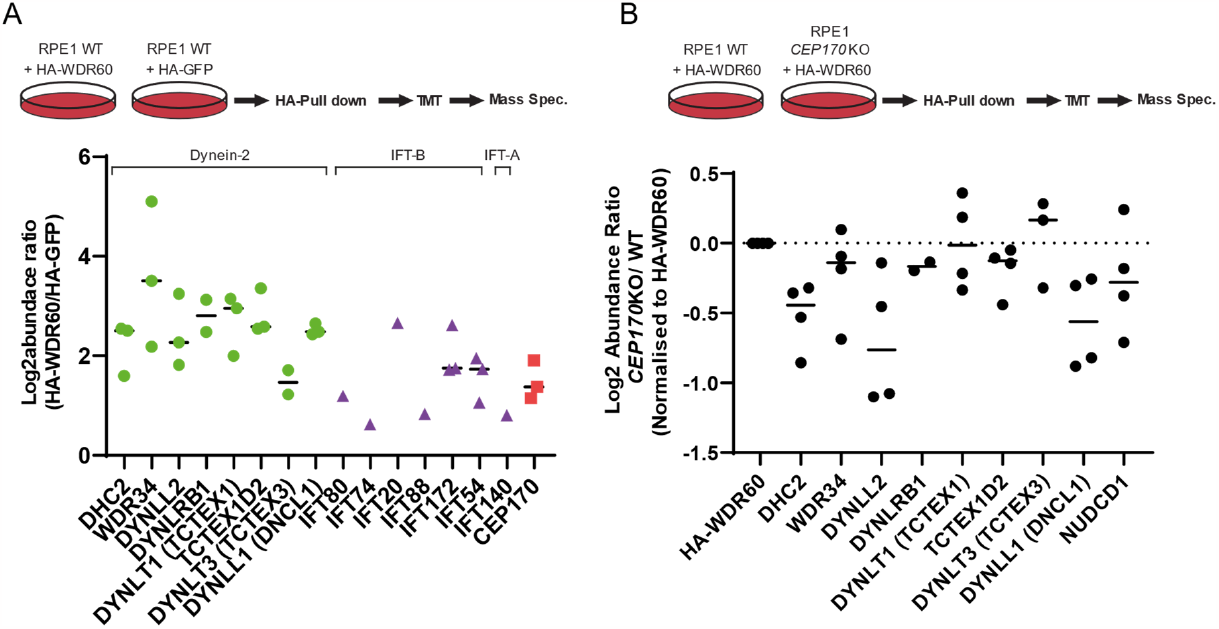
Disruption of the dynein-2 holocomplex in CEP170 KO cells. TMT-proteomics from RPE1 WT cells and *CEP170* KO (clones 24H8 and 23G7). Cells were transfected with HA-GFP and HA-WDR60 (A) or HA-WDR60 (B). (A) Dynein-2 and IFT components and CEP170 are detectable in RPE1 cells (dynein-2 subunits, green circles, IFT subunits, purple triangles, CEP170). We did not detect CEP170B in this dataset (see also Table S1). (B). HA-WDR60 interaction with dynein-2 components is decreased (Log_2_ relative abundance <0) in *CEP170* KO cells. Three independent experiments were performed. Data are normalized to HA-WDR60 levels. Bars represent means.

## Discussion

A functional cilium requires an intricate assembly of many multi-protein complexes including dynein-2, IFT-A, and IFT-B. Our data show that CEP170 interacts with dynein-2 and promotes its assembly. Loss of CEP170 does not prevent ciliation or cause significant defects in cilia function, suggesting a possible role for CEP170 as a modulator of cilia. CEP170 was originally identified as a binding partner of polo-like kinase 1 (PLK1) in a yeast two-hybrid screen and shown to localise to sDAPs on mother centrioles (Guarguaglini et al., 2005). From this initial work it was clear that CEP170 has a role in microtubule organisation and subsequent studies have also linked it with mitosis and DNA damage responses (Qin et al., 2019; Rodriguez-Real et al., 2023; Welburn and Cheeseman, 2012; Zhang et al., 2019). CEP170 has also been shown to interact with the TZ protein RPGRIP1L (Gupta et al., 2015). CEP170 may act as a hub, connecting the cell cycle and cilia disassembly with dynein-2 and cilia function.

### CEP170 and sDAPs in ciliogenesis and cilia function

The mother centriole is distinct from the daughter owing to the addition of DAPs and sDAPs. The precise components of these structures are not fully defined, with new partners being identified and better microscopy techniques continuing to enable better definition of their exact location on centrioles. DAPs contain a core set of proteins, including ODF2 (Ma et al., 2023). sDAPs are comprised of ODF2, CEP128, centriolin, CCDC120, CCDC68, ninein, α- and γ-taxilin, and CEP170 (Ma et al., 2023). The consensus is that whilst DAPs are required for ciliogenesis, sDAPs are mostly seen to be redundant and are more important for microtubule anchoring (Hall and Hehnly, 2021; Mazo et al., 2016). Indeed, there are currently no reported ciliopathies associated with any known sDAP gene. Loss of ODF2 has been shown to affect ciliogenesis, but whether this is from their role in the sDAPs is not clear as ODF2 is also present in DAPs (Ishikawa et al., 2005; Kashihara et al., 2019). CEP170 is located at the tip (distal) point of sDAPs, but additional non-sDAP centrosomal localisation has been observed (Guarguaglini et al., 2005; Kashihara et al., 2019; Mazo et al., 2016). Previous studies showed that depleting either CCDC120, CCDC68, or ninein is sufficient to remove CEP170 from sDAPs, and that this did not affect ciliation (Huang et al., 2017; Mazo et al., 2016). Our results are broadly consistent with these observations, cells lacking CEP170 were still able to generate cilia of comparable lengths to WT cells. Whilst we observed normal length cilia in IMCD3 *CEP170* KO cells, we do note that they ciliated to a lower degree, suggesting a more general role for CEP170 and sDAPs in cilia spatial control as previously observed (Mazo et al., 2016). However, we also see defects in cilia function, not previously studied in this context. Takahara et al. (2018) reported that whilst KO of the IFT-A component, IFT122, led to severe ciliogenesis defects, KO of other IFT-A genes had minor effect on ciliogenesis but with impaired trafficking in ciliary proteins (Hirano et al., 2017; Takahara et al., 2018). In *CEP170* KO cells, whilst we observed near-normal IFT velocities (Lechtreck, 2015), we did see accumulation of IFT88 at the ciliary tip, as well as impacts on SAG-dependent Smo-localisation.

Cilia assembly and disassembly is linked to the cell cycle, only non-dividing cells are capable of primary cilium assembly (Breslow and Holland, 2019; Ishikawa and Marshall, 2011; Mirvis et al., 2018). Accordingly, during cell cycle progression the primary cilium must be disassembled. The processes that govern this are not fully defined and multiple mechanisms have been suggested (Breslow and Holland, 2019; Mirvis et al., 2018; Phua et al., 2017; Pugacheva et al., 2007). In our study, *CEP170* KO cells had delayed cilia disassembly, as shown previously using depletion of *CEP170* by RNA interference (Lamla, 2009). In contrast, in *WDR60* KO cells cilia are disassembled at a rate comparable to WT cells, suggesting that the delay occurs independently to dynein-2. CEP170 may, therefore, act as a point of integration between the cell cycle and cilia disassembly, possibly via already known interactions between CEP170 and the microtubule depolymerising kinesins, KIF2A/B (Maliga et al., 2013; Miyamoto et al., 2015; Welburn and Cheeseman, 2012).

### CEP170 and dynein-2 assembly

Along with cilia defects following loss of CEP170, the main conclusion of this work is the identification of, CEP170, as an interactor of dynein-2. To date, the majority of published dynein-2 interactions have only been reported in the context of IFT-A and IFT-B (Hiyamizu et al., 2023a; Hiyamizu et al., 2023b; Shak et al., 2023; Toropova et al., 2019; Tsurumi et al., 2019; Vuolo et al., 2018; Zhu et al., 2021) these interactions have given us great insight into how dynein-2 interacts with IFT-A and IFT-B. Whilst we cannot say for certain which subunit of dynein-2 CEP170 binds to, CEP170 can be robustly detected using either WDR34 and WDR60 in immunoprecipitations. Immunoprecipitation of HA-WDR60 from *CEP170* KO cells compared to WT cells reveals roles for CEP170 in dynein-2 holocomplex assembly and in maintaining DHC2 localization at the basal body. Given the reduction in dynein-2 holocomplex it might be expected that the cilia defects would be much more severe. However, in *Chlamydomonas*, depletion of endogenous DHC2 by approximately 50% does not result in any drastic defects in cilia (Reck et al., 2016). Even when DHC2 levels were more substantially reduced, cilia were still able to form. Moreover, loss of *WDR60* in nematodes and in RPE1 cells, does not ablate cilia formation or function (De-Castro et al., 2022; Vuolo et al., 2018), suggesting that cilia require only small amounts of dynein-2 holocomplex to function. This raises questions as to why cells maintain a larger pool of dynein-2 than is needed to maintain cilia and IFT.

Dynein-2 light, light-intermediate and intermediate chains (WDR34 and WDR60) are required for dynein-2 assembly (Vuolo et al., 2018), stabilise dimerization of the heavy chain, and support formation of the auto-inhibited state (Perrone et al., 2003; Rompolas et al., 2007; Toropova et al., 2019). In this study, we find that immunoprecipitation of HA-WDR34 from *WDR60* KO cells captures more CEP170 than from WT cells. Considering these data together, it is possible that CEP170 interacts with intermediate chains during dynein-2 assembly and/or stabilises the complete holocomplex. CEP170 could also act in capturing dynein-2 as retrograde trains disassemble on exit from cilia (van den Hoek et al., 2022). In these ways, CEP170 could serve to enrich dynein-2 at sDAPs to promote IFT train assembly. The KIF3A subunit of the anterograde IFT motor kinesin-2 also localizes to sDAPs (Kodani et al., 2013), perhaps indicating a broader role for sDAPs acting as a “shunting yard” in recruitment of components for eventual IFT train assembly.

## Materials and Methods

Unless stated otherwise, all reagents were purchased from Sigma-Aldrich (Poole, UK).

### Plasmids

pLVX plasmids encoding HA-WDR60, HA-WDR34, HA-GFP and GFP-Arl13b were generated previously (Vuolo et al., 2018).

### Cell culture

Human telomerase-immortalized retinal pigment epithelial cells (hTERT-RPE1, ATCC CRL-4000) were grown in DMEM-F12 (Gibco) supplemented with 10% fetal bovine serum (FBS) (Gibco) at 37°C with 5% CO_2_ . Cells were not validated further after purchase from ATCC. *WDR60* KO RPE1 cells were generated previously (Vuolo et al., 2018). Mouse inner medullary collecting duct (IMCD-3, ATCC CRL-2123) and IMCD3-*FlpIn-IFT88-NG3* (a gift from M. Nachury, UCSF) cells were grown in DMEM/F-12(HEPES) (Gibco) supplemented with 5% FBS, 100 U/ml penicillin-streptomycin at 37°C with 5% CO_2_.

### Ciliogenesis and cilia disassembly

RPE1 and IMCD3 cells were washed twice in phosphate-buffered saline (PBS) and incubated in serum-free medium for 24 hrs to induce ciliogenesis. For Smo experiments, confluent cells were placed in serum-free media and treated with Shh agonist SAG (Selleckchem (from Stratech Scientific, Ely, UK) Catalog No.S7779) at the final concentration of 400 nM for 24 hrs. A cilium disassembly assay was performed as described in Zhang et al. (2019). Specifically, cells were starved in serum-free medium for 48 hrs to induce cilium formation. Serum was then added back to the medium to stimulate cilium resorption. Cells were harvested at various time points for immunolabelling assays. To monitor cilia excision, RPE1 WT and *CEP170* KO cells (clone 24H8) stably expressing GFP-Arl13B, were grown on imaging dishes and serum starved. Serum was added 30 minutes before imaging at 37°C on inverted an inverted fluoresence Leica-TIRF microscope using a 60×/1.40 oil objective. Images were collected every 2-4 minutes until the cilia became out of focus or photobleached.

### Genome editing

The guide RNAs (gRNA) targeting human *CEP170* (RPE1 cells), mouse *CEP170* and mouse *CEP170B* (IMCD3 cells) were designed using ‘chop chop’ software (Labun et al., 2016).

#### RPE1 cells

The gRNA sequences (5’-CAACTATGATGCGTCTA-3’; and 5’-TGGGCAGCCGTCATCGT -3’) were designed to target exon 3 and exon 7 of human *CEP170*. TrueCut Cas9 Protein v2 (Invitrogen) and TrueGuide Synthetic gRNA targeting *CEP170* gene locus (CRISPR1123823_CR and CRISPR1123824_CR, Invitrogen) were cotransfected with TrueCut™ Cas9 Protein v2 and crRNA+tracrRNA (7.5 pmol each). into cells using Lipofectamine CRISPRMAX (Invitrogen) according to the manufacturer’s instructions. After 48 hrs, cells were sorted, and single cells were plated in a 96-well plate (Corning). To check the *CEP170* gene, genomic DNA was extracted and the target sequences subjected to PCR to amplify targetted exons. The PCR products were cloned in the pGEM T Easy vector system (Promega) according to the manufacturer’s instructions and sequenced (Eurofins Genomics and SourceBioscience). One clone was generated with with exon 3 edited (*CEP170* KO 23G7) and one with exon 7 edited (*CEP170* KO 24H8). Small deletions causing a frameshift were detected in both alleles (Fig. S2A and B). *CEP170* KO was confirmed by immunoblotting with RPE1 WT cells (Fig. S2C).

#### IMCD3 cells

The gRNA (5’-CGAGAAATGATTTTCGT-3’) was designed to target exon 2 of mouse *CEP170*. Similarly, the gRNA (5’-CCACGAAGATGAGTTCACG-3’) was designed to target exon 2 of mouse *CEP170B*. pSpCas9(BB)-2A-GFP (Addgene plasmid, #PX458) was used as the vector to generate a gRNA of both *CEP170* and *CEP170B*. In a well of a 6-well plate (Corning), 50–60% confluent IMCD3 cells were transfected with 2.4 μg plasmid DNA, using 8.5 μl Lipofectamine 2000 transfection reagent (Thermo Fisher Scientific) in 300 μl Opti-MEM reduced serum medium (Gibco). After 48 hrs, GFP-positive cells were sorted and cultured for two weeks before single cell sorting. Single cell sorted cells were seeded in 96-well plates (Corning) in 150 μl conditioned medium. Conditioned medium was prepared by sterile filtration of a 1:1 mixture of fresh medium, and medium removed from a flask of cells in exponential growth phase (50–70% confluent). Single cell sorted colonies were expanded by subculture to increasingly larger culture vessels (24-well, 12-well plates, T25 flask). *CEP170* and *CEP170B* gene editing was confirmed as described for RPE1 cells. One clone was validated for *CEP170* KO (*CEP170* KO A2G7), with a small deletion detected in each allele causing a frameshift (Fig S2D), validated by immunoblot (Fig. S2E). One clone was validated for *CEP170B* KO (*CEP170B* KO B2A11),with deletions detected in each allele causing a frameshift (Fig S2F), validated by immunoblot (Fig. S2G). For IMCD3-*FlpIn-IFT88-NG3* cells, two clones (both with exon 2 edited) were validated (*CEP170* KO A2B10 and *CEP170* KO A2A5) with changes to exon 2 (Fig. S6). Clone A2B10 has an 8 base pair deletion in one allele and a large insertion in the second allele (Fig S6A). Clone A2A5 had deletions detected in both alleles, one causing a frameshift (Fig S6B). Both clones were validated by immunoblot (Fig. S6C).

### Antibodies

The antibodies used, and their dilutions for immunoblotting (IB) and immunofluorescence (IF) are as follows: mouse anti-acetylated tubulin (Sigma T6793, IF 1:2000, lot number 0000108922), rabbit anti-IFT88 (Proteintech 13967-1-AP, IF 1:300, lot 00044070), rabbit anti-TMEM67 (MKS3) (Proteintech 13975-1-AP, 1:50 IF, lot 00022584), rabbit anti-RPGRIP1L (Proteintech 55160-1-AP, IF 1:100, lot 09000273), rabbit anti-DHC2 (DYNC2HC1) (ab122525, IB 1:200, lot GR247356-8), rabbit anti-DHC2 (DYNC2HC1) (ab225946, IF 1:50, lot GR3220321-11), rabbit anti-Arl13B (Proteintech 17711-1AP, IF 1:1000, lot 00076202), mouse anti-Smo (Santa Cruz sc-166685, IF 1:100, lot E1721), mouse anti-GAPDH (HRP-6004, lot 21002053, 1:1000 IB), rabbit anti-CEP170 (proteintech 27325-1-1, IF 1:500, IB 1:1000, lot 00054066), rabbit anti-CEP170B (Sigma HPA000871, IF 1:250, IB 1:300, lot r95478) mouse anti gamma-tubulin (Sigma t5326, IF 1:500, lot 10m4782v), rabbit anti-WDR34 (Novus NBP188805, 1:300 IB, lot A105337), rabbit anti-LIC3 (Proteintech 15949–1-AP, IB 1:250, lot 00007524), mouse anti-Ninein (Proteintech, 67132-1-Ig, IF 1:500, lot 10008943). AlexaFluor-conjugated secondary antibodies were from Invitrogen (A21206, A10037, A10042, A21202, IF 1:500), HRP-conjugated secondary antibodies were from Jackson ImmunoResearch (115-035-166, 111-035-144, IB 1:10,000).

### Immunofluorescence

Cells grown on 0.17 mm thick (#1.5) coverslips (Fisher Scientific, Loughborough, UK) were washed twice in PBS, and then fixed in ice-cold methanol at −20°C for 5 minutes. For Smo labelling, cells were fixed for 10 minutes at room temperature (RT) in 4% paraformaldehyde (PFA) and permeabilized with PBS containing 0.1% Triton X-100 for 5 minutes. For DHC2 (DYNC2HC1) labelling, cells were permeabilized with PBS containing 0.3% Triton X-100 for 10 minutes. Subsequently, cells were blocked with 3% bovine serum albumin (BSA) in PBS for 30 minutes. The coverslips were incubated with primary antibodies for one hour, washed in PBS and then incubated with relevent secondary antibodies for one hour. Nuclear staining was performed using 4,6-diamidino-2-phenylindole (DAPI) (Life Technologies) at a concentration of 1 μg·ml^-1^ in PBS for 3 minutes. Cells were then rinsed twice in PBS before mounting on glass slides (VWR) using Mowiol 4-88 mounting medium. Cells were imaged using an Olympus IX-71 widefield microscope with a 63x oil immersion objective (N.A. 1.4), and excitation and emission filter sets (Semrock, Rochester, NY) controlled by Volocity software (version 4.3, Perkin-Elmer, Seer Green, UK). For ciliary DYNC2HC1 (Fig. 5A and B), cells were imaged on a Leica SP5 confocal microscope system (Leica Microsystems, Milton Keynes, UK). Images were acquired as 0.5 μm z-stacks.

#### Immunoblotting

Cells were lysed in ice-cold buffer containing 50 mM tris(hydroxymethyl)aminomethane hydrochloride (Tris-HCl) pH 7.5, 150 mM NaCl, 1% Igepal and 1 mM ethylenediaminetetraacetic (EDTA) with protease inhibitor cocktail (539137, Millipore). Samples were separated by SDS-PAGE followed by transfer to nitrocellulose membranes (Cyvita). Membranes were blocked in 5% milk in tris-buffered saline with 0.1 % tween-20 (TBST). Primary antibodies diluted in 5% milk-TBST were incubated with membrane overnight at 4°C. Membranes were washed in TBST, incubated with secondary antibodies for 1 hour, washed with TBST and detected by enhanced chemiluminescence (Promega).

### Fluorescence intensity measurements

Quantification of fluorescence intensity was performed using average z-stack projections of original images in ImageJ (Schindelin et al., 2012). Local background normalised fluorescence intensity was measured at the cilary base or along the axoneme using the plot profile tool after manually tracing the axoneme in the cilary marker channel.

### Immunoprecipitation

Anti-HA immunoprecipitation was performed as previously described (Vuolo et al., 2018).

### TMT-labelling and proteomic analysis

Proteins on HA-agroase beads were digested with trypsin, TMT-labelled and analysed by mass spectometry as previously described (Vuolo et al., 2018). Three independent experiments were performed with data displayed as normalised Log_2_ abundance ratios.

### Live-cell IFT88 TIRF imaging

Cells were seeded (1x10^5^) on a 35 mm glass bottom imaging dish (MatTek) in normal media (DMEM/F12, HEPES with 5% FBS, Gibco). The day after, cells were washed twice in PBS before being serum starved in phenol red-free media (DMEM/F12, HEPES, Gibco) for 24 hours. Cells were imaged at 37°C with CO_2_ on an Olympus/Abbelight SAFe360 system with two Hamamatsu Fusion sCMOS cameras, 488 nm diode laser and 100x oil immersion objective. The pixel size was 99.7 nm. Cell surface plane was found using automated TIRF angles in the mNeonGreen channel. Abbelight NEO software was used for acquisition of movies, with frames captured every 100 ms for no longer than 1 minute per movie and no more than 1 hour per dish.

### IFT88 TIRF kymograph analysis

Movies of individual cilia were imported into ImageJ (Schindelin et al., 2012). Each cilium was manually traced and kymographs (in both directions) were generated using the KymographClear plug-in for ImageJ (http://www.nat.vu.nl/~erwinp/downloads.html, Prevo et al. (2015)). Individual events were manually traced, and the gradient was converted to a speed (μm·s^-1^) by dividing by frame length (100 ms) and multiplying by pixel size (99.7 nm). The average velocity from each cilium was plotted. As we cannot tell the tip from the base, anterograde IFT was defined as being the faster of either direction. Where no obvious trace could be unambiguously traced, the velocity was called as 0 μm·s^-1^.

### Electron microscopy

Cells were grown on 35 mm dishes (Corning) before being fixed in 2.5% glutaraldehyde for 20 minutes and processed as previously described in Vuolo et al. (2018). Sections were imaged on an FEI (Cambridge, UK) Tecnai12 transmission electron microscope.

## Acknowledgements

We thank members of the Stephens lab for helpful discussion and advice, especially M. Esther Prada-Sanchez for technical assistance. The IMCD3-*FlpIn-IFT88-NG3* cells were a gift from Max Nachury, UCSF. The authors gratefully acknowledge the University of Bristol Wolfson Bioimaging Facility and Flow Cytometry Facility for their support and assistance in this work.

## Author contributions

Conceptualization: A.J.R., D.J.S.; L.V.

Methodology: J.F.W., L.V., C.S., A.G.M., L.R.H., and K.H.

Validation: J.F.W., L.V., C.S., and L.R.H.

Formal analysis: J.F.W., L.V., C.S., D.J.S. and L.R.H.

Investigation: J.F.W., L.V., A.P., C.S., K.H. and L.R.H.

Data curation: J.F.W., L.V., and D.J.S.

Writing - original draft: J.F.W., L.V., D.J.S.

Writing - review & editing: J.F.W., L.V., C.S., A.G.M., L.R.H., A.J.R., and D.J.S.

Visualization: J.F.W., L.V., C.S., and L.R.H.

Supervision: J.F.W., L.V., D.J.S.; A.J.R.

Project administration: D.J.S.; A.J.R.

Funding acquisition: D.J.S.; A.J.R., L.V.

## Data availability

The mass spectrometry proteomics data have been deposited to the ProteomeXchange Consortium via the PRIDE partner repository with the dataset identifier PXD046827 and PXD046844.

