## Supplementary Figures and Legends for "Roles for CEP170 in cilia function and dynein-2 assembly": Supplementary Material biorxiv.pdf

| Bait | $Log_2$ abundance ratio | | Reference | EBI PRIDE Accession Code[1] |
|---|---|---|---|---|
| | CEP170 | CEP170B | | |
| **TMT** | | | | |
| **GFP-WDR34** | 1.161 | Not found | Shak et al., 2023 | PXD032758 |
| | 1.347 | Not found | Shak et al., 2023 | PXD032758 |
| | 1.310 | Not found | Shak et al., 2023 | PXD032758 |
| | 2.139 | 1.266 | n/a | PXD046827 |
| | 2.160 | Not found | n/a | PXD046827 |
| | 1.643 | Not found | n/a | PXD046827 |
| **HA-WDR34** | 2.522 | Not found | Hiyamizu et al., 2023b | PXD031151 |
| | 1.854 | 0.908 | Vuolo et al., 2018 | PXD010398 |
| **HA-WDR60** | 1.919 | Not found | Hiyamizu et al., 2023b | PXD031151 |
| | 1.110 | 2.143 | Hiyamizu et al., 2023b | PXD031152 |
| | 1.910 | 0.915 | Hiyamizu et al., 2023b | PXD031153 |
| | 0.633 | 0.174 | Hiyamizu et al., 2023b | PXD031154 |
| | 1.907 | Not found | This paper (Fig. 7A) | PXD046844 |
| | 1.375 | Not found | This paper (Fig. 7A) | PXD046844 |
| | 1.147 | Not found | This paper (Fig. 7A) | PXD046844 |
| **Non-TMT[2]** | | | | |
| **GFP-WDR34** | Found | Not found | Hiyamizu et al., 2023b | PXD031157 |
| | Found | Found | Hiyamizu et al., 2023b | PXD031158 |
| | Found | Not found | Hiyamizu et al., 2023b | PXD031156 |
| **GFP-WDR60** | Found | Found | Hiyamizu et al., 2023b | PXD031157 |
| | Found | Found | Hiyamizu et al., 2023b | PXD031158 |
| | Found | Not found | Hiyamizu et al., 2023b | PXD031156 |

**Table S1**: Summary table of CEP170 or CEP170B found in previous dynein-2 interaction proteomics. [1]EBI Proteomics IDEntifications (PRIDE) database (https://www.ebi.ac.uk/pride/, Perez-Riverol et al. (2022)). [2]These data sets were not isobarically labelled and so we do not report relative abundances.

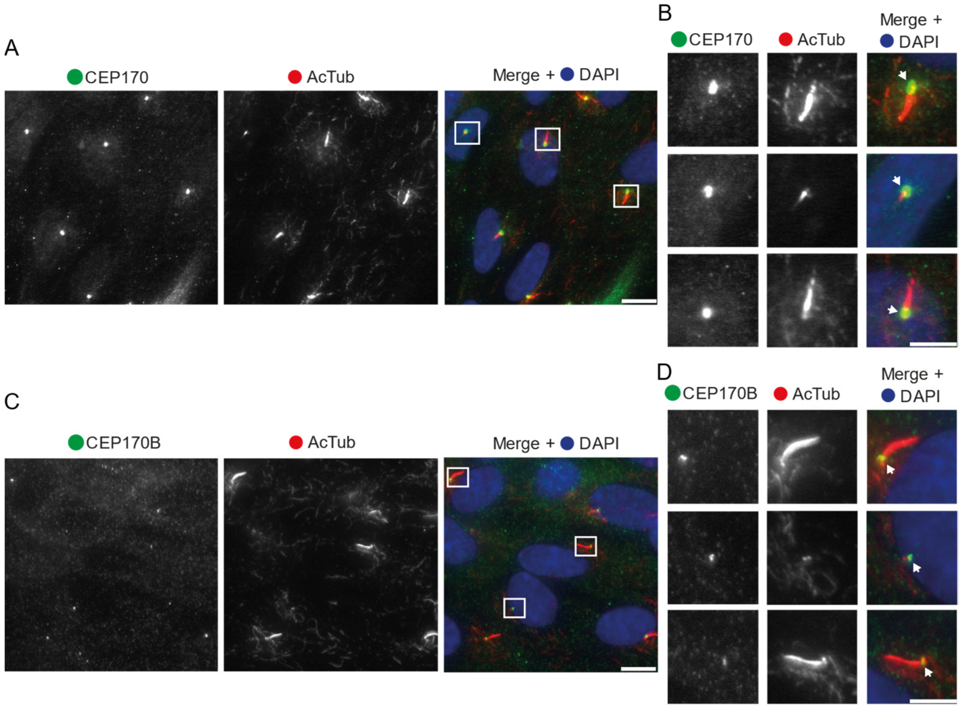


***Figure S1: Centrosomal localisation of CEP170 and CEP170B***. RPE1 WT cells were serum starved and fixed. Cells were stained for either CEP170 (A and B) or CEP170B (C and D) (green). Cilia were labelled with acetylated tubulin (AcTub, red). B and D are zoom of regions from A and C, respectively. Scale bars = 5 µm. Arrows indicate the centrosome.

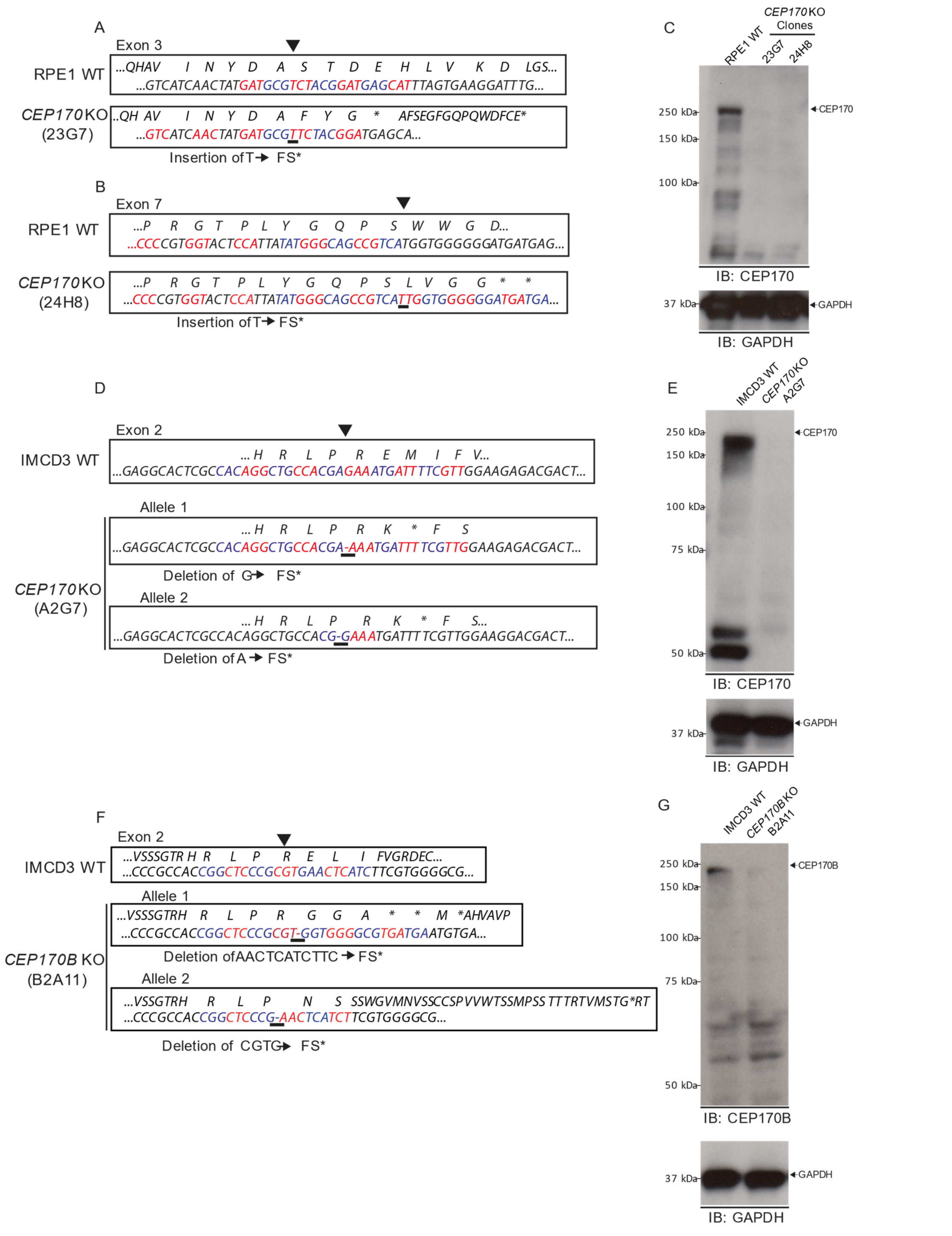


***Figure S2: Mutation analysis for RPE1 and IMCD3 KO cell lines***. (A) Alignment of human *CEP170* exon 3 indicating edited region for *CEP170* KO (clone 23G7) RPE1 cells. A single base pair (bp) insertion causes a frameshift and premature stop codon (FS*). (B) Alignment of human *CEP170* exon 7 indicating edited region for *CEP170* KO (clone 24H8) RPE1 cells. A single bp insertion causes a frameshift and premature stop codon (FS*). (C) Immunoblot (IB) on whole cell lysates from *CEP170* KO clones 23G7 and 24H8 showing CEP170 is no longer present. GAPDH is shown as loading control. (D) Alignment of mouse *CEP170* exon 2 indicating edited region for *CEP170* KO (clone A2G7) IMCD3 cells. Different mutations were found on each allele, each a single bp deletion causing a frameshift-stop. (E) Immunoblot showing absence of CEP170 in clone A2G7. GAPDH is shown as a loading control. (F) Alignment of mouse *CEP170B* exon 2 indicating edited region for *CEP170B* KO (clone B2A11) IMCD3 cells. Different mutations were found on each allele. Allele 1 had an 11 bp deletion, allele 2 had a 4 bp deletion, each causing a frameshift-stop. (G)

Immunoblot showing absence of CEP170B in clone B2A11. GAPDH is shown as a loading control. Arrowheads above alignments indicate point of modification (underlined).

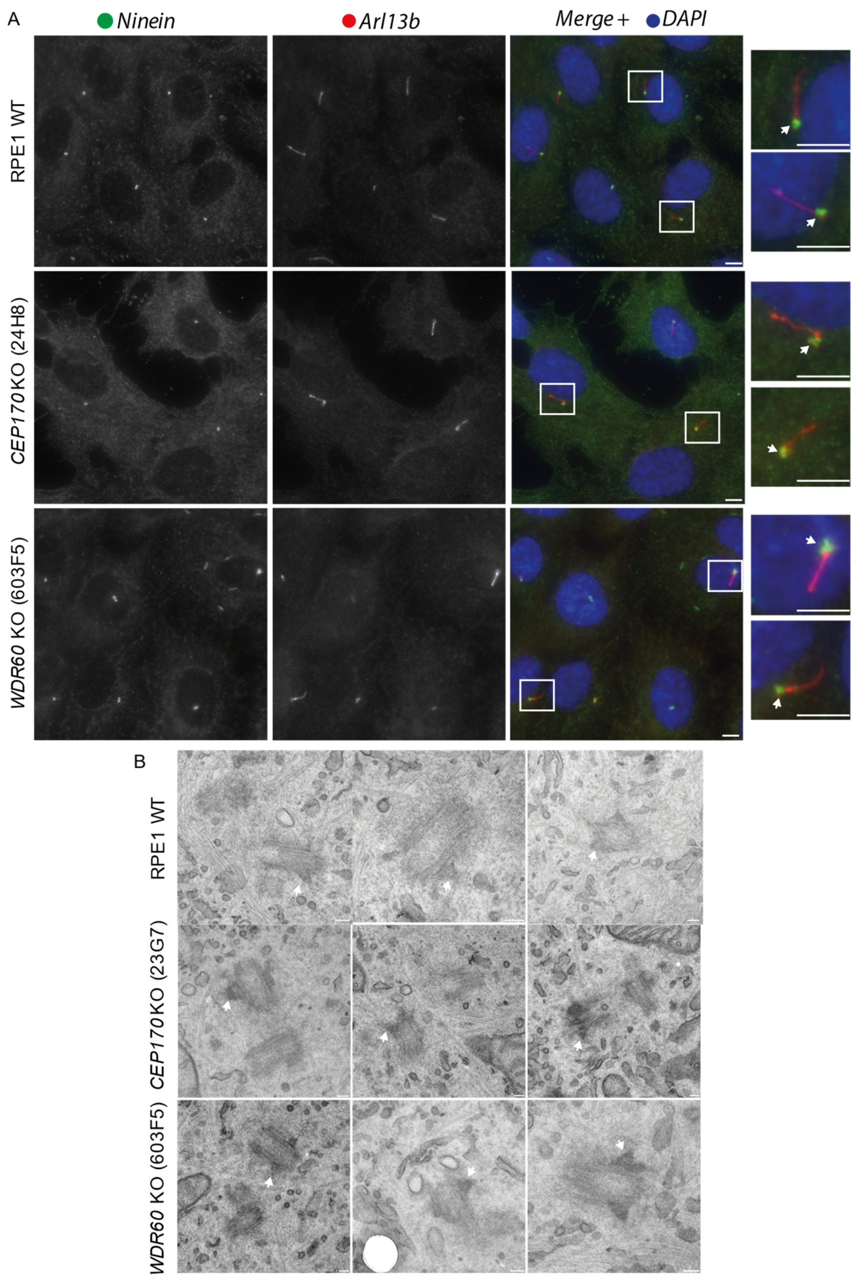


***Figure S3: Normal sDAPs in CEP170 KO cells.*** (A) Ciliated WT RPE1, *CEP170* KO (clone 24H8) and *WDR60* CKO (clone 603F5) cells were stained for ninein (green) and the cilia marker Arl13B (red). Ninein localises to the basal body (indicated with arrows). Scale bar = 5µm. (B) EM of non-ciliated WT RPE1, *CEP170* KO (clone 23G7) cells to examine the centrioles. sDAPs can be seen on the mother centriole (indicated with arrows). Three examples are shown in each case. Scale bars = 100 nm.

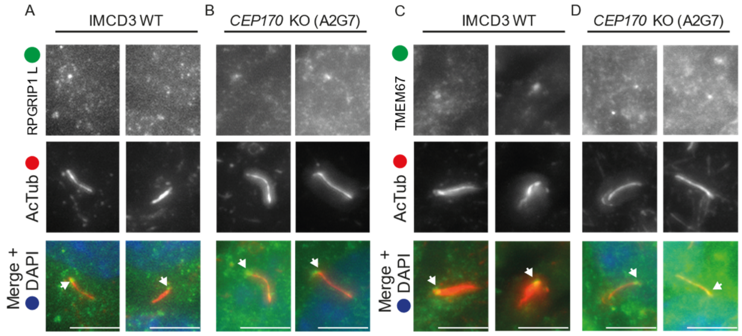


***Figure S4: TZ is unaffected in CEP170 KO IMCD3 cells***. IMCD3 WT and *CEP170* KO (clone A2G7) cells were grown, serum starved and fixed before staining for TZ markers (green), RPGRIP1L (A+B) or TMEM67 (C+D). Cilia were labelled with acetylated tubulin (AcTub, red). Arrows indicate TZ. Scale bars = 5 µm.

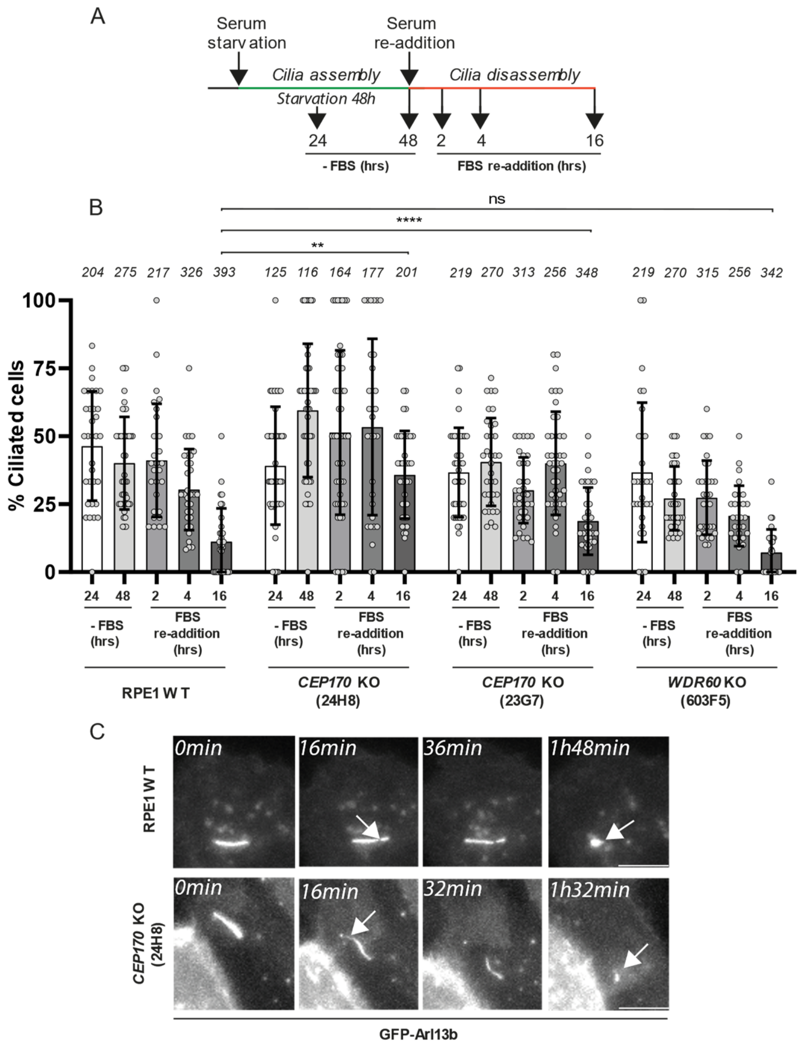


***Figure S5: Cilia disassembly following serum re-addition***. RPE1 WT, *WDR60* KO (603F5) and *CEP170* KO (clones 24H8 and 23G7) cell lines and serum starved for 48 hours before serum was re-added and cells incubated for indicated time points (shown in A) before fixing. Cells were stained for ciliary markers (Arl13b/AcTub, as in Fig. 2) and cilia number (%) was quantified (B). Data is represented as mean ± SD. For each cell line at least three independent experiments were carried out. For each time point, the total number of cells counted is shown above in italics. Mann-Whitney two-tailed test was used to detect statistical significance at 16hrs after re-addition of serum, ** – p=0.0047, **** – p<0.0001. (C)

Live imaging of ciliated RPE1 WT or *CEP170* KO cells stably expressing GFP-Arl13b. Serum was re-added 30 minutes prior to imaging. Arrows indicate cilium excision points. Scale bar = 5 µm.

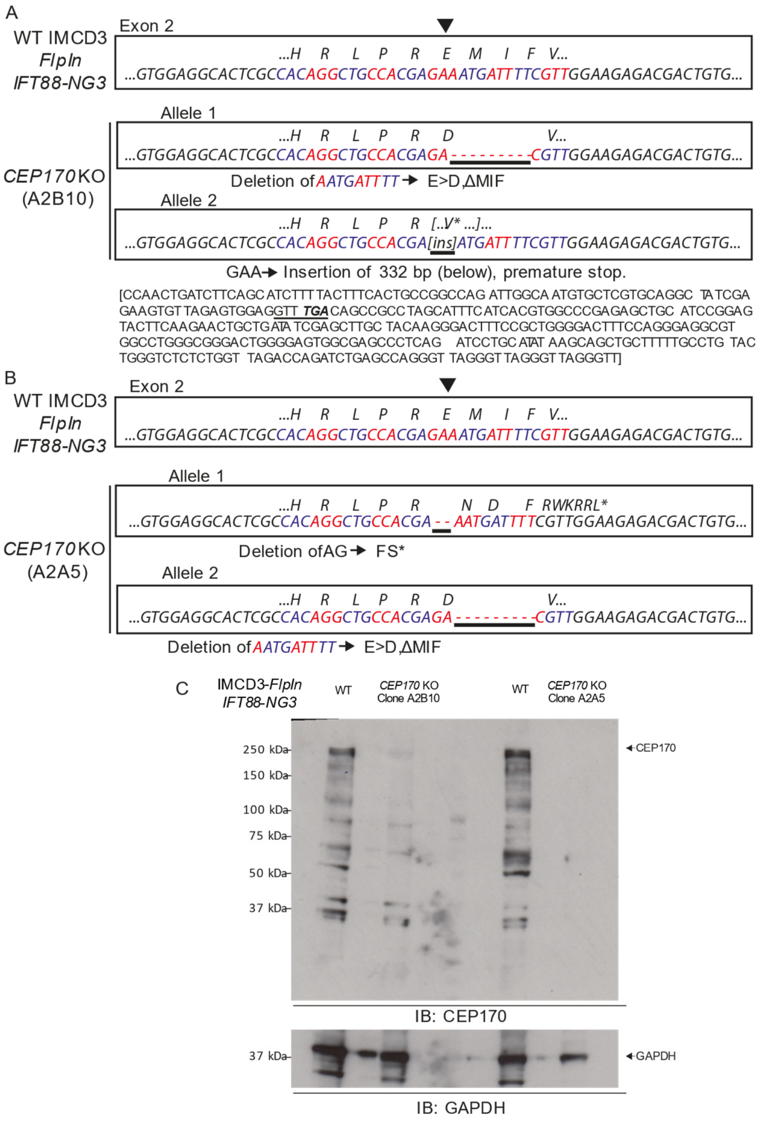


***Figure S6: Mutation analysis for IMCD3 FlpIn-IFT88-NG3 CEP170 KO cell lines.*** (A) Alignment of mouse *CEP170* exon 2 indicating edited region for *CEP170* KO (clone A2B10) IMCD3 *FlipIn-IFT88-NG3* cells. Different mutations were found on each allele. Alleles 1 had a 9 bp deletion, causing a Glu to Asp change, and triplet amino acid deletion (ΔMet-Ile-Phe). Allele 2 had a 3 bp substitution for a 332 bp region with a downstream stop codon (underlined, in-frame stop codon (TGA) is in bold italics). (B) Alignment of mouse *CEP170* exon 2 indicating edited region for *CEP170* KO (clone A2A5) IMCD3 *FlpIn-IFT88-NG* cells. Different mutations were found on each allele. Allele 1 had a 2 bp deletion causing a frameshift-stop. Allele 2 had a 9 bp deletion, causing a Glu to Asp change, and triplet amino acid deletion (ΔMet-Ile-Phe). Arrowheads above alignments indicate point of modification (underlined). (C) Immunoblot (IB) on whole cell lysates from *CEP170* KO clones A2B10 and A2A5 showing CEP170 is no longer present. GAPDH is shown as loading control.

**Movies 1-3: Representative movies from live-cell TIRF imaging**. Relates to Fig. 4. (1) IMCD3 *FlipIn-IFT88-NG3*, (2) *CEP170* KO (clone A2B10), (3) *CEP170* KO (clone A2B5). Time is indicated in each movie; playback is in real-time. Scale bars = 2 µm.
